# Water transmission potential of *Angiostrongylus cantonensis*: larval viability and effectiveness of rainwater catchment sediment filters

**DOI:** 10.1101/496273

**Authors:** Kathleen Howe, Lisa Kaluna, Bruce Torres Fisher, Yaeko Tagami, Robert McHugh, Susan Jarvi

**Author notes:** These authors contributed equally to this work (Kathleen Howe, Lisa Kaluna).

## Abstract

Neuroangiostrongyliasis, caused by *Angiostrongylus cantonensis*, has been reported in Hawai‘i since the 1950’s. An increase in cases is being reported primarily from East Hawai‘i Island, correlated with the introduction of the semi-slug *Parmarion martensi.* Households in areas lacking infrastructure for water must use rainwater catchment as their primary domestic water supply, for which there is no federal, state, or county regulation. Despite evidence that contaminated water can cause infection, regulatory bodies have not addressed this potential transmission route. This study evaluates: 1) the emergence of live, infective-stage *A. cantonensis* larvae from drowned, non-native, pestiforous gastropods; 2) larvae location in an undisturbed water column; 3) longevity of free-living larvae in water; and 4) effectiveness of rainwater catchment filters in blocking infective-stage larvae. Larvae were shed from minced and whole gastropods drowned in either municipal water or rainwater with >94% of larvae recovered from the bottom of the water column. Infective-stage larvae were active for 21 days in municipal water. Histological sectioning of *P. martensi* showed proximity of nematode larvae to the body wall of the gastropod, consistent with the potential for shedding of larvae in slime. Gastropod tissue squashes showed effectivity as a quick screening method. Live, infective-stage larvae were able to traverse rainwater catchment polypropylene sediment filters of 20 μm, 10 μm, 5 μm, and 1 μm filtration ratings, but not a 5 μm carbon block filter. These results demonstrate that live, infective-stage *A. cantonensis* larvae can and do emerge from drowned snails and slugs, survive for extended periods of time in water, and that the potential exists that they enter the household water supply. This study illustrates the need to better investigate and understand the potential role of contaminated water as a transmission route for neuroangiostrongyliasis.

## Introduction

The nematode *Angiostrongylus cantonensis* is established throughout the main Hawaiian Islands with the possible exception of Lāna‘i [1, 2, 3]. The complex lifecycle of this parasite has been well-described in the literature [4, 5, 6, 7]. In Hawai‘i, *Rattus rattus* and *Rattus exulans* are important definitive hosts, and many gastropod species are effective intermediate hosts including *Achatina fulica*, *Euglandia rosea*, *Laevicaulis alte*, *Limax maximus*, *Parmarion martensi* and *Veronicella cubensis* [1, 2, 8]. The third stage larva (L3) is harbored in the intermediate host, and it is this larval stage that is infective to rats and accidental hosts, including humans, as the L3 larvae can safely pass through the acidic environment of the mammalian gut. There are also paratenic hosts that can carry the infective stage larvae; these include shrimp, prawns, crabs, frogs, water monitor lizards, centipedes, and some planarians [7. 9, 10, 11, 12]. Of planarians, the predacious *Platydemous manokwari*, the New Guinea flatworm, has been determined to be an important carrier of *A. cantonensis* [9].

Infection by *A. cantonensis* is reported as the leading cause of eosinophilic meningitis (EM) worldwide [10, 13, 14]. As the parasite targets the central nervous system, the disease can cause serious and irreparable harm. The first cases of neuroangiostrongyliasis, or rat lungworm disease, were reported in 1959 on O‘ahu, and both victims died as a result of infection [15]. A review of medical cases of EM in the State of Hawai‘i from 2001-2005 identified 83 cases of meningitis, 24 of which were attributed to neuroangiostrongyliasis [13]. Hawai‘i Department of Health (HDOH) reports cluster cases began to occur on Hawai‘i Island in 2004-2005, and there has been a steadily increasing trend of severe cases, with 107 cases of neuroangiostrongyliasis from 2001-2017. Of these, 77 have originated from Hawai‘i Island [16].

While human infection on Hawai‘i Island has been attributed to “lifestyle” [17], the trend of case increases actually correlates with the introduction of an effective intermediate host, *P. martensi* the semi slug, which was first reported on O‘ahu in 1996 and on Hawai‘i Island in 2004 [8]. A survey conducted in 2005 found a 77.5% infection rate in this species [8] in East Hawai‘i. While many species can be intermediate hosts, *P. martensi* harbors higher parasite loads compared with other hosts [2, 18, 19]. Quantification by real-time PCR of *P. martensi* tissue samples collected in Hawai‘i in 2005 determined an average burden of 445 larvae per 25 mg tissue versus 1-250 for other gastropod species. In 17% of *P. martensi* collected, real-time PCR results showed burdens of more than 1000 larvae per 25 mg tissue [19]. At these concentrations, it would seem that ingestation of even a small piece of tissue could cause a serious case of disease. Also, *P. martensi* exhibits unusual behavior in that it is relatively fast, has a propensity to climb, and is attracted to human dwellings and food items [8]. The increase in cases of neuroangiostrongyliasis on Maui [16] may be related to the establishment of *P. martensi* populations which have been anecdotally reported on Maui by an author of this paper (K.H.) and have been substantiated by Cowie et al. [20].

Disease transmission is generally thought to occur from intentional or accidental ingestion of infected intermediate or paratenic host organisms on unwashed or poorly washed produce or from undercooked hosts [10, 16]. Some patients believe they were infected through exposure to contaminated rainwater catchment. The use of rainwater catchment as a source of household and/or agricultural water is prevalent on East Hawai‘i Island, where most cases of neuroangiostrongyliasis originate. While exact numbers are not known, it was estimated that 30,000-60,000 people relied on catchment water when the Guidelines on Rainwater Catchment Systems for Hawai‘i manual was written in 2010 [21]. In the Puna District, where most recent cases have originated [16, 22, 23], there are large subdivisions which were developed in the late 1950’s to mid 1970’s with little or no infrastructure for water [24]. Today, the majority of households in this district rely on rainwater catchment for their household water supply and there is no state or federal agency that oversees the use or management of catchment systems [25]. The design, installation, and maintenance of rainwater catchment systems can be expensive and laborious and can create systems that do not provide potable water at all taps, which is required of public water systems. Hawai‘i homeowners with mortgages from the Veterans Affairs are required to have a copy of the “Guidelines on Rainwater Catchment Systems for Hawai‘i,” which makes recommendations for roofing, gutters, tanks, tank covers, sediment filters and water treatments [21]. HDOH recommends but does not require homeowners to implement the guidelines. Contractors or local vendors can also provide guidance regarding system design, installation, and maintenance but this information may not be consistent among providers. A recent survey conducted on Hawai‘i Island showed 90% of respondents used their catchment water for drinking or bathing, but that only 66% of these respondents had catchment systems that might be expected to provide water safe to drink [25]. Many residents and catchment tank cleaners report finding slugs and snails in catchment tanks, likely seeking access for moisture or having been washed down rain gutters (Fig 1).

**Fig 1.**
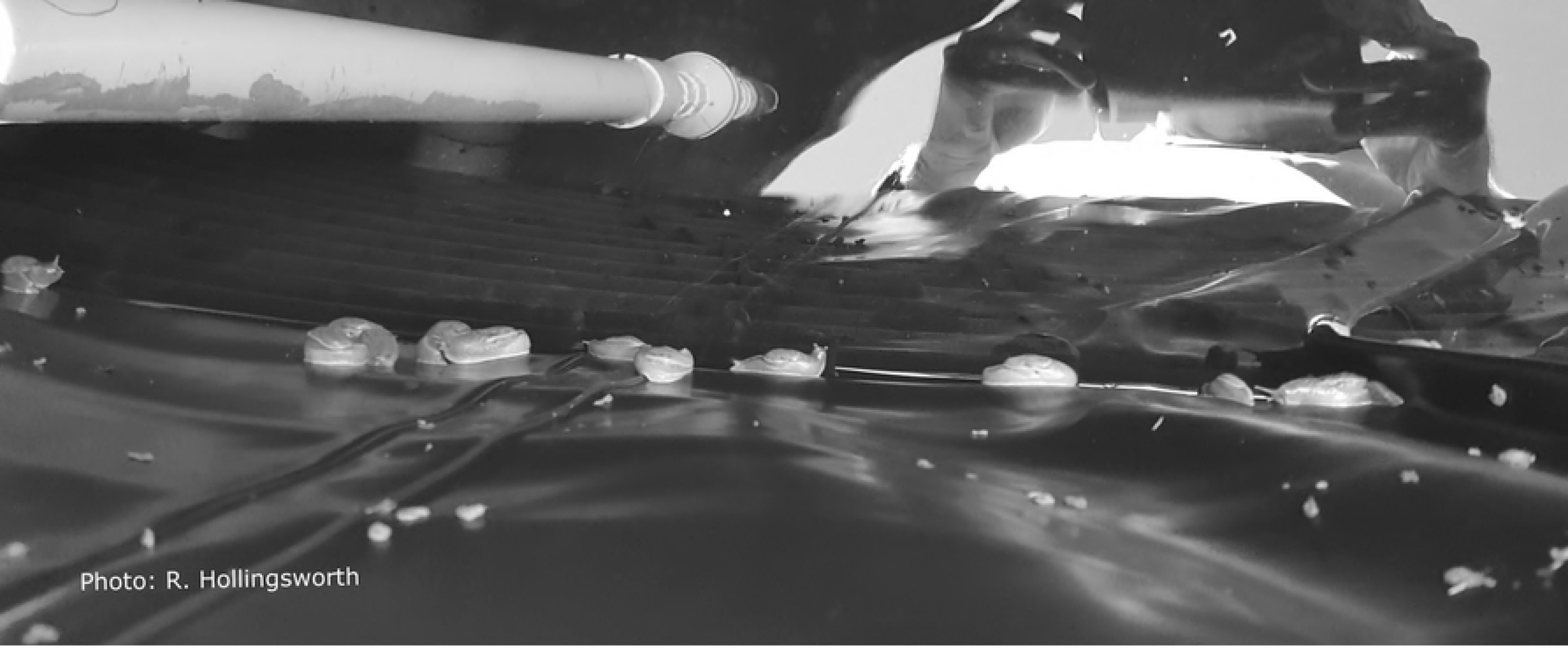
Slugs inside rainwater catchment tank. Looking into a rainwater catchment tank with many *P. martensi* on the plastic liner just above the water line (reflection in water, water intake pipe, and pulled back cover shown). The tank was tightly covered, however the slugs were still able to access the tank. Photo credit; R. Hollingsworth.

Ash [26] examined the morphology of infective-stage (L3) *A. cantonensis* (n = 35) and determined a mean width of 26 μm with a range of 23-34 μm. The 2010 Rainwater Catchment Guide states that a 20 μm catchment sediment filter should be sufficient to prevent passage of the larvae; however, no formal filter studies have been conducted in Hawai‘i. Moreover, many manufacturers attest that their sediment filters will only “reduce” number of microorganisms and that their micron sizing is based on nominal particulate ratings of >85% of a given size as determined from single-pass particle counting results [27]. This rating system does not take into account microorganism behavior and/or their ability to burrow through or swim around a filter when the pump system is turned off. Early studies confirm L3 larvae shed from drowned or live gastropods in water were subsequently infective to rats. Cheng and Alicata [28] demonstrated that both uninjured and intentionally injured *A. fulica*, *Subulina octona*, and *L. alte* shed larvae when partially submerged in municipal water. Uninjured snails shed fewer L3 larvae (2-10) than injured snails (55 L3 larvae). Larvae survived for up to 72 hours and when fed to rats were recovered as young *A. cantonensis* adults after 17 days. Richards and Merritt [29] confirmed these findings, showing larvae shed from snails into fresh water were active for at least seven days, and that rats became infected after drinking water containing L3 larvae. A third study by Crook, Fulton, and Supanwong [30] described *A. fulicia* crawling into wells and water jars, and well water contamination with *A. cantonensis* was reported. Their study showed that of 30 *A. fulica* drowned in sedimentation funnels, 18 were infected and shed larvae which were used to successfully infected *Rattus norvegicus.* If rats can be infected by drinking L3 contaminated water it is possible that humans and other mammals may also be infected in this manner. Currently, no studies have been conducted in Hawai‘i to determine the larval shedding potential of the efficient, recently introduced intermediate host *P. martensi*.

The relationship between the widespread use of rainwater catchment and/or exposure to contaminated water sources, the introduction of the effective intermediate host *P. martensi*, and the high incidence of neuroangiostrongyliasis on Hawai‘i Island may be of epidemiological significance. Therefore, this study was conducted to evaluate the larval shedding potential of drowned gastropods, particularly of *P. martensi*, and to assess larval longevity in water. A pilot study was also initiated to evaluate the effectiveness of commercially available sediment filters in reducing or blocking *A. cantonensis* larvae in a laboratory-based model catchment system.

## Materials and methods

### Gastropod collection and preparation

Slugs and snails used in all studies were non-native considered invasive species. Specimens were collected in the the Koa‘e and Wa‘a Wa‘a area in the lower Puna District of Hawai‘i Island and in the nearby Hilo District, both areas of known *A. cantonensis* infection. Collection sites in Puna were on private land and approximately a three linear-mile distance from each other (Fig 2). In Hilo, collecting was done on the campus of the University of Hawai‘i. Captured gastropods were held in individual collection tubes or bags to avoid cross-contamination. Species collected included *A. fulica*, *L. alte*, *P. martensi*, and *V. cubensis*; however, *P. martensi* was the primary gastropod of interest. Tissue tail snips were excised and weighed from all gastropods except where noted. Tissue samples were used for tissue squashes or placed in 100 μL DNA lysis buffer (0.1M Tris HCl, 0.1M EDTA, 2% SDS) for subsequent genetic analysis. The remainder of the gastropod was placed in a 50 mL falcon tube filled with rainwater or municipal water, and inverted until the gastropod was deceased.

**Fig 2.**
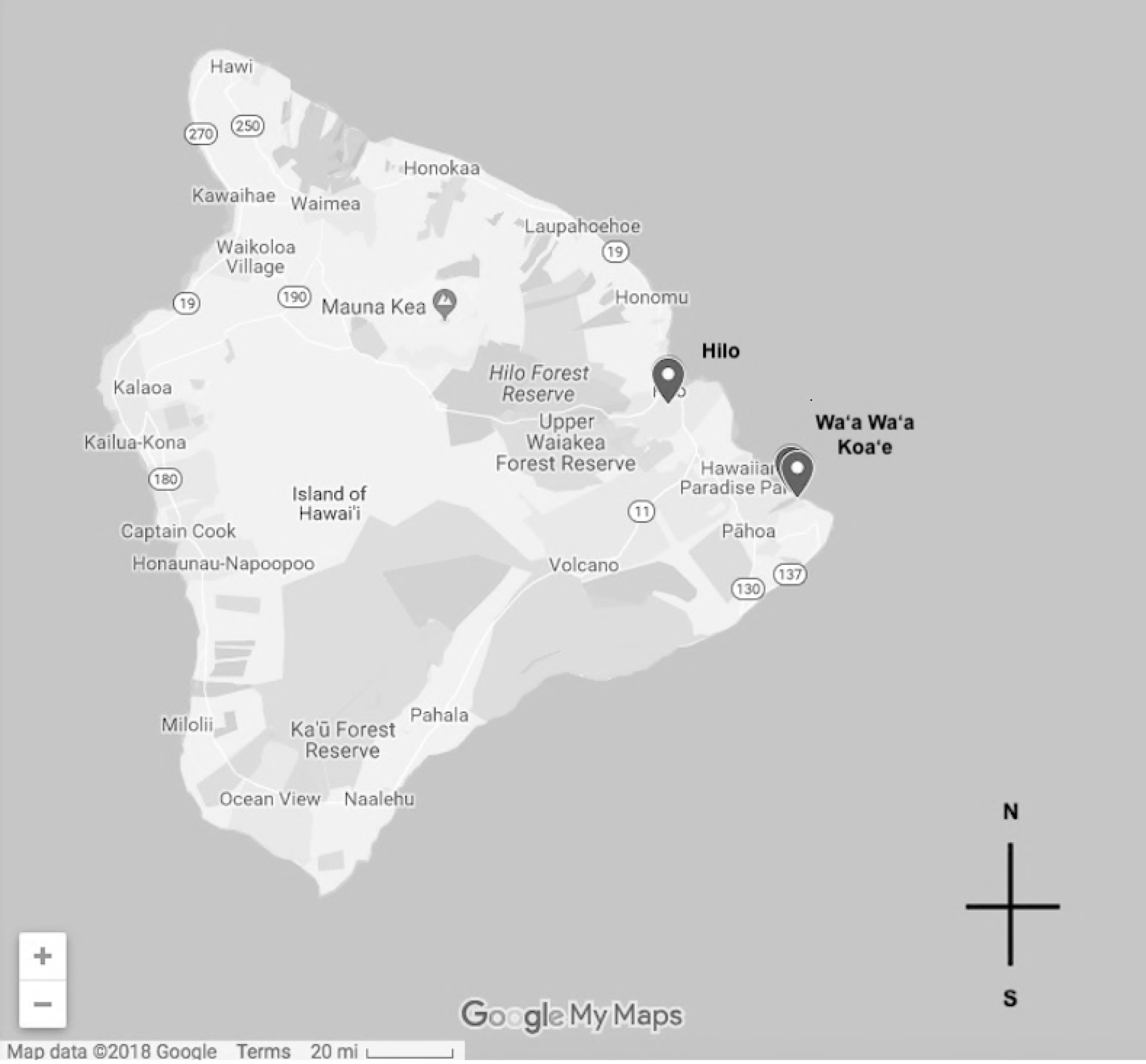
Map of gastropod collection sites. Gastropods were collected on the east end of Hawaii Island in the Wa‘a Wa‘a and Koa‘e area, and in Hilo, on the University of Hawaii campus.

### Potential for shedding of *A. cantonensis* larvae by drowned gastropods

#### Rainwater collection

Rainwater was collected in a clean, food-grade, ~19 L bucket placed below the drip edge of a clean, gutter-less metal roof and was transfered into clean two-liter glass jars. Control samples of water (15 mLs) were pipetted into sterile 60mmX 15 mm disposable petri dishes and regularly observed by microscopy for evidence of larvae using a dissecting microscope (Leica EZ4, Wetzlar, Germany). Samples were labeled and held at room temperature (~21° C) and were repeatedly inspected over the course of the trial for evidence of emerged larvae. A 200μL sample of the rainwater was tested with real-time PCR (see below) for presence/absence of the parasite [18, 31].

#### Larval location in a water column

Water from gastropods drowned for whole vs minced and diversified gastropod species experiments was used to evaluate the distribution of larvae in a water column after a gastropod drowning event. Water samples (5 mL) were pipetted from three locations in tubes containing a gastropod drowned in rainwater: the top, the middle, and the bottom (TMB) of the 50 mL water column. Each 5 mL sample was placed into a petri dish and 10 mL of additional water added to the dish to prevent drying. After sampling, each Falcon tube was topped off with rainwater. The first samples were drawn within 24 hours post drowning (PD). Samples were then taken at 24-hour intervals for as few as five, and as many as 20 days PD. Petri dishes were held at room temperature (~ 21° C) and were routinely examined (24-72 hours) for evidence of larvae by microscopy. Larvae were counted, photographed, and isolated for genetic analysis. All photography of larvae was done using an Olympus CX31 compound microscope and LW Scientific MiniVID USB 5MP Digital Eyepiece Camera (Lawrenceville, GA) and ToupView v. 3.7.

#### Whole versus mechanically minced *P. martensi*

As the literature suggested that damaged gastropods shed more larvae, a trial was conducted using *P. martensi* to determine if this species could shed larvae when drowned, and if damage had an effect on larval shedding. Ten *P. martensi* slugs were collected and processed as described above, and then randomly assigned to a treatment group (whole n = 5, or minced n ; 5). Slugs were mechanically minced with single-use safety blades and placed in 50 mL rainwater and whole slugs were drowned in rainwater. Samples were taken from the TMB at 24-hour intervals over a 96-hour timeframe. Petri dishes were examined daily for ten consecutive days for the quantity of larvae. Two-sample t-tests (Minitab 18) were used to evaluate the difference in mean larval loads, as determined by qPCR, the difference in slug weights, and differences in numbers of larvae shed between whole versus minced slugs.

#### Diversified gastropod species

Larval shedding was evaluated across multiple gastropod species including *A. fulica* (n = 4), *L. alte* (n = 2), *V. cubensis* (n = 2), and *P. martensi* (n = 4). Gastropods were collected, species were equally divided into treatment groups (whole or minced) and processed as described above in rainwater. Water samples (5 mLs) were taken from the TMB for examination of larvae, with sampling beginning at day 0, and samples taken every 24 hours for 20 days with volumes replaced daily.

#### Sieve separation of varied-size larvae and longevity trials

Ash [26] reported L3 larvae diameters of 23-34 μm, which is close to the 20 μm sediment filter size recommendation from the Guidelines on Rainwater Catchment Systems in Hawai‘i [21]. Trials were conducted to separate and identify larvae shed from drowned, mechanically damaged gastropods (*L. alte* = 2, *P. martensi* = 2). Shed, active larvae of various sizes, which were never observed to be coiled in form, were challenged to traverse a 20 μm metal sieve (Hogentogler & Co., Columbia, Maryland). The sieve was seated in a beaker with a volume of municipal water covering the top of the mesh and larvae were pipetted onto the sieve. The sieve was removed after 24 hours and the liquid below was examined by microscopy. Larvae found below the sieve were removed from the beaker by pipette and were held in petri dishes. A subsample of these larvae was placed into acid (0.5% HCl/0.5% pepsin) to observe larval reaction [2]. The remaining larvae (~1000) were held to determine longevity. Subsamples of these larvae (~250) were processed for genetic analysis at 53 and 56 days PD. Shed larvae, which were initially observed to be coiled and had emerged into active larvae, were also challenged to traverse the sieve. Subsamples were exposed to acid and were held for observation for longevity. At 21 days PD ~80 larvae were isolated for genetic analysis. C-shaped larvae were not used in sieve trials as these larvae were never observed to be active. The sieve was soaked in a 15% salt solution for 20 minutes, rinsed in soapy water followed by a fresh water rinse, dried at ~ 50° C, and exposed to 2000 Joules of UV radiation (UVP CX-2000 UV Crosslinker, Upland, CA) to destroy any DNA between trials.

#### Municipal water versus rainwater

*P. martensi* (n = 16) were used to determine if water source had an effect on larval shedding. Tail snips were taken from 10 slugs for subsequent real-time PCR testing and six were left whole with no tail snips taken. The slugs were divided into two treatments: 50 mL of either municipal water or rainwater (tail snip slugs n = 5 per group, and whole slugs n = 3 per group). Three 5 mL samples were drawn from the bottom of tubes at 24, 48, 72, and 96 hours PD and placed into individual petri dishes. All samples drawn were examined daily and larvae were counted and isolated for genetic analysis.

### Tissue squash to screen for presence of larvae

To determine the effectiveness of tissue squashes as a screening method for nematode infection in slugs, a small piece of tail tissue (~5 mg) was removed from the tail snip of *P. martensi* (n = 10) for evaluation. The remaining tail snip was used for genetic analysis. Tissue was placed between two glass slides and pressure was applied until a thin film was achieved. The slide was examined with an Olympus CX31 compound microscope for visualization of larvae.

### Histology

Several *P. martensi* were prepared using traditional histological methods to examine location of larvae in the tissue [32]. The shells were first removed and gastropods were immersed in glacial acetic acid for 24 hours to dissolve any remaining shell fragments. The specimens were fixed in 10% formalin for 48 hours and then transferred to 70% ethanol, after which they were cut laterally into three sections (head, middle, tail). The sections were processed in a tissue processor (Leica TP 1020, Leica Microsystems Inc., Bannockburn, IL), blocked in wax, cut in 7μm sections which were placed on glass slides, and stained with traditional hematoxylin and eosin. Slides were examined with an Olympus CX31 compound microscope. Sections containing larvae were photographed as described above.

### Catchment Sediment Filter Testing

A laboratory-based catchment system (Fig 3) was constructed replicating a home design common in East Hawai‘i dwellings, with the exception of the size of the water reservoir and the absence of a separate pressurized tank [21]. All components were approved for potable water use. A 132 L pressure tank (Sta-rite SR35-10S) was used as a water reservoir, filled with municipal water, and connected directly to a water pump (Grundfos 96860195). Polyvinyl chloride (PVC) piping in 3/4″ diameter (JM Eagle 57471) connected all components of the system. PVC primer and cement (Oatey 30756, and 31013) was used to seal all non-threaded connections and plumbers’ tape (Oatey 0178502) was used to seal all threaded connections. A pressure gauge (ASME B40.1:1991) was installed just prior to the nematode loading station to ensure pressure during testing replicated a home environment. A one-way check valve (ProLine 101-604HC) was installed just prior to the nematode loading station, to prevent backflow of experimental nematodes into the system. Two PVC ball valves (ProLine 107-634HC) cut off water flow just prior to the filter housing, in between these, a tee socket fitted with a socket male adapter and threaded cap (Charlotte 187917, 188131, and 536725) allowed for loading of experimental nematodes for each trial. Five commercially available sediment filters, obtained from local vendors, were tested inside a universal housing (Pentair 158215). Filtrate was directed into a 19 L water bottle (ORE International WS50GH-48) through vinyl tubing (Watts 032888192362). The mouth of the water bottle was fitted with flexible PVC coupling (Fernco 687960) and a PVC ball valve (ProLine 107-634HC) to use as a spigot.

**Fig 3.**
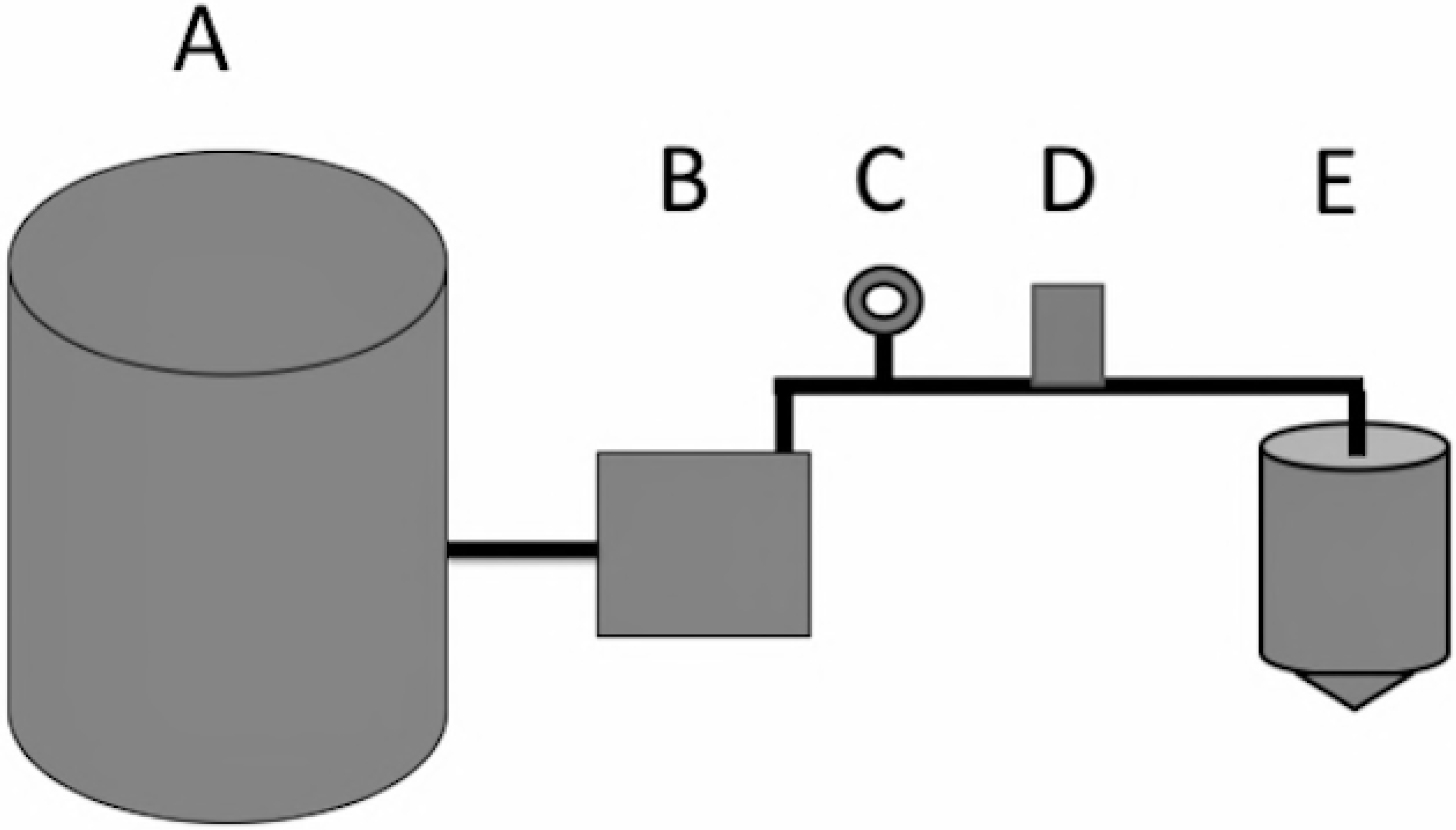
Design of the laboratory-based catchment system. Basic layout of the model catchment system used, with a (A) 132-liter water reservoir, (B) a water pump, (C) a pressure gauge, (D) a nematode loading station, (E) universal housing with sediment filter, and (F) a 19-liter collection tank with a spigot all connected with ¾″ PVC piping. (Figure is not accurately scaled).

For each trial, live nematodes were individually isolated into fresh municipal water from whole, intact *P. martensi* collected in the Hilo area and drowned in a 50 mL Falcon tube of municipal water as described above. Prior to loading into the system, nematodes were visualized on a stereoscope (Wild Heerbrugg M5A APO) where ~16% of nematodes were photographed at 50X total magnification, using a microscope digital camera (MiniVID TP605100) and software (ToupView 3.7). Length and diameter of nematodes were measured using ToupView 3.7 following calibration with a stage micrometer (American Optical 1400). Width was measured at the widest point of the nematode, as determined by visual analysis. Nematodes were added to the loading station as described above. The water pump was run for approximately 10 seconds, yielding roughly 10 L of filtrate in the collection tank. The filtrate was immediately transferred into 1-liter bottles and then vacuum filtered across a 0.2 μm nylon filter (Whatman 7402-004) to concentrate and isolate post-filter nematodes. Nematodes on the surface of the 0.2 μm filter were rinsed off using a wash bottle of municipal water and observed by microscopy. Resulting nematodes were counted, observed for movement, and up to 25 nematodes per filter were measured and isolated for genetic analysis. The nematodes isolated for genetic analysis were kpooled per individual filter. Measurements of post-filter nematodes for the first replicate of the 1 μm spun polypropylene filter were not obtained due to an inability to obtain high resolution images of decomposing nematodes from a six-day delay in observing the vacuum filtrate. The proportion of pre- and post-filter nematodes that were infective *A. cantonensis* L3 larvae was calculated by comparing nematode length to that previously reported [26]; width would be an inappropriate comparison as the location of measurement on the nematode was nonspecific. Morphological analysis was used in conjunction with genetic analysis to determine whether infective *A. cantonensis* L3 larvae were among the post-filter nematodes for each filter replicate.

New filters of each type of sediment filter were tested in either duplicate or triplicate. Each individual filter was left in the system continuously for four runs of 250 nematodes loaded in runs 1 and 2, and 500 nematodes loaded in run 3. Time between runs 1-3 was contingent upon nematode availability and varied from 15 minutes to 16 days. To see if previously introduced nematodes could live in or on the filter and penetrate it over time, the system was run a fourth time, seven or eight days after run 3, without loading additional nematodes. Following the fourth run of a filter, the catchment system was disinfected by allowing a 10% bleach solution to completely fill the system and collection tank for 20 minutes. Subsequently, the entire system was thoroughly rinsed with municipal water by running nearly 390 L of water through the system (3 volumes of the water reservoir) before a new filter was installed. Each new filter was flushed with municipal water for 15 minutes before testing. Nematode loss from the system and vacuum filtration process was independently measured to establish recovery rates. To test nematode loss from the entire testing process, excluding the catchment sediment filter, three runs of 250 nematodes each were tested in the system (including vacuum filtration) with no sediment filter inside the housing. To test nematode loss attributable to the vacuum filtration process, 100 nematodes were added to just the vacuum filtration apparatus and counted three times, post-vacuum filtration.

Data analysis: As not all data were normally distributed, non-parametric Kruskal-Wallis tests were used to test for significant differences between the proportions of post-filter nematodes between filter type and between runs within each filter test. Non-parametric Wilcoxan tests were used to examine the differences in the number of post-filter nematodes between replicates of the 20 μm and 10 μm filters, and a Kruskal-Wallis test for the 5 μm spun replicates. Statistical analysis was conducted in Minitab 18.

### Development of Cloned Reference Standards

Plasmids used for standards and positive controls were made by cloning the amplicon of an *A. cantonensis* larval sample used in Jarvi et al. [18]. The target region was amplified from genomic DNA using real-time PCR as described above. The amplification product was removed from a 2% low-melt agarose gel and purified using a DNA, RNA, and protein purification kit (Macherey-Nagel 740609.240C) following the manufacturers protocol. The purified product was cloned using the Invitrogen TOPO TA Cloning Kit for Sequencing (Thermo Fisher Scientific K4575J10) and colonies were screened for the target insert by PCR, all per manufacturer’s protocol. Colonies with the target insert were grown overnight at 37° C in 7 mL of TYE medium containing 50 μg/mL ampicillin, and plasmids were isolated using a QIAprep Spin Miniprep Kit (Qiagen 27104) per the manufacturer’s protocol. Plasmids were sequenced with M13F/M13R primers on an Applied Biosystems 3730XL DNA Analyzer at the Advanced Studies in Genomics, Proteomics and Bioinformatics Sequencing Facility at the University of Hawai‘i at Manoa. The sequences were verified using the GenBank BLAST analysis. One plasmid was chosen for use in real-time and quantitative PCR. Eight serial dilutions (1:10 - 1:10^8^) of the chosen plasmid were made using Buffer AE (Qiagen); 1-2 μl were quantified by qPCR using the same methods and genomic DNA described in Jarvi et al. [18] and analyzed as described above. All samples, standards, and non-template controls were run in duplicate. After analyzing the standard curve, the mean quantities (# larvae per reaction) of the eight plasmid dilutions were used in the qPCR run testing the gastropods described below.

### qPCR of gastropod tissue

DNA extractions were completed using a DNeasy Blood & Tissue Kit (Qiagen 69506) per the manufacturer’s Animal Tissue protocol with a few adjustments. Gastropod tissues in 100 μl DNA lysis buffer were digested with 180 μl of buffer ATL and 20 μl of proteinase K, ending with a final elution of 400 μl. DNA was quantified using a Bio-Spec Nano (Shimadzu Scientific Instruments Inc., Carlsbad California). Extracted DNA was subjected to qPCR using species-specific primers that were redesigned into a custom assay [18, 31]. Samples were run in triplicate. Reactions were run on a StepOne Plus RealTime PCR system (Life Technologies, Carlsbad CA) with minimum modifications to the assay manufacturer’s cycling conditions (1 cycle of 50°C for 2 min, 95°C for 10 min, followed by 40 cycles of 95°C for 15 sec, 60°C for 1 min). Optical tubes (Life Technologies 4358297) were exposed to ten minutes of ultra-violet radiation (UVC, 254 nm) and reactions were 20 μl in volume with either 100 ng of total DNA per reaction or the maximum allowed template volume of 9 μl.

Data analysis: StepOne Software v2.3 was used for analysis of all runs using the auto-threshold setting and verifying all replicates of non-template controls had no exponential amplification before data was used. Samples and standards were determined positive if all replicates showed exponential amplification in both the ‘Δ Rn vs Cycle’ and ‘Rn vs Cycle’ plot types with a cycle threshold standard deviation (CTSD) of <0.5. The number of larvae per mg of tissue in the tail snips was determined as follows:

\# of larvae/mg = (# larvae per reaction/ template vol (μl)) × Final elution vol (400 μl)/ tail snip weight (mg)

### Real-time PCR of larvae

Larvae shed from gastropods were collected by pipette for real-time PCR analysis into DNA lysis buffer, allowed to settle, and supernatant was removed leaving ~100 μl of liquid. The concentrated larvae were homogenized in a glass tissue grinder for approximately five minutes and DNA was extracted as described above. The tissue grinder was cleaned with a 10% bleach solution and thoroughly rinsed between uses. Post-filter nematodes from the catchment sediment filter trials were isolated by pipette and DNA was extracted as above with a final elution of 50 μl. Larvae samples were run in either duplicate or triplicate with positive controls of plasmid standards for the determination of the presence of *A. cantonensis* DNA.

## Results

### Potential for shedding of *A. cantonensis* larvae by drowned gastropods

#### Rainwater

At no time were larvae or other live organisms observed by microscopy in the 10 mL samples of clean rainwater collected. The real-time PCR result for the 200 μL rainwater sample was negative for *A. cantonensis*.

#### *P. martensi* (whole vs minced)

Larvae were shed from both whole (n = 5) and minced (n = 5) *P. martensi* that were drowned in rainwater, and qPCR results of all tail snips were positive for *A. cantonensis* (Table1). There was no significant difference in weight between the whole and minced groups (P = 0.586). Quantification of *A. cantonensis* in the tail snips ranged from 4.62-39.20 larvae per milligram of tissue. Cycle threshold (C_T_) values of the standards ranged from 16-32 cycles and the unknown samples were 20-26 cycles, all with CT SD < 0.5. The standard curve had an R^2^ value of 0.961, a slope of −3.677, a y-intercept of 22.862, and a PCR efficiency of 87.051%. The larvae load averages between treatment groups was not significant (P =0.590), but the numbers of total larvae shed between treatment groups was significant (P = 0.043) with greater numbers shed by whole *P. martensi.* Noticeable larval shedding occurred at 72 and 96-hours post-drowning (PD). Shed larvae were observed to be either coiled or C-shaped and inactive. The coiled larvae were observed to emerge from this state to become vigorously swimming larvae, while the C-shaped larvae exhibited no movement or emergence. The C-shaped larvae fit the description of L2 larvae as described by Lv et al. [33]. The active larvae had the morphological features of the L3 *A. cantonensis* and displayed the characteristic S and Q-movement described for *A. cantonensis* and not displayed in other free-living nematode species [33]. At ten days PD > 200 actively swimming larvae remained in petri dish samples. Of the three sampling locations, the bottom samples contained 93.5% of all larvae shed.

**Table 1.**
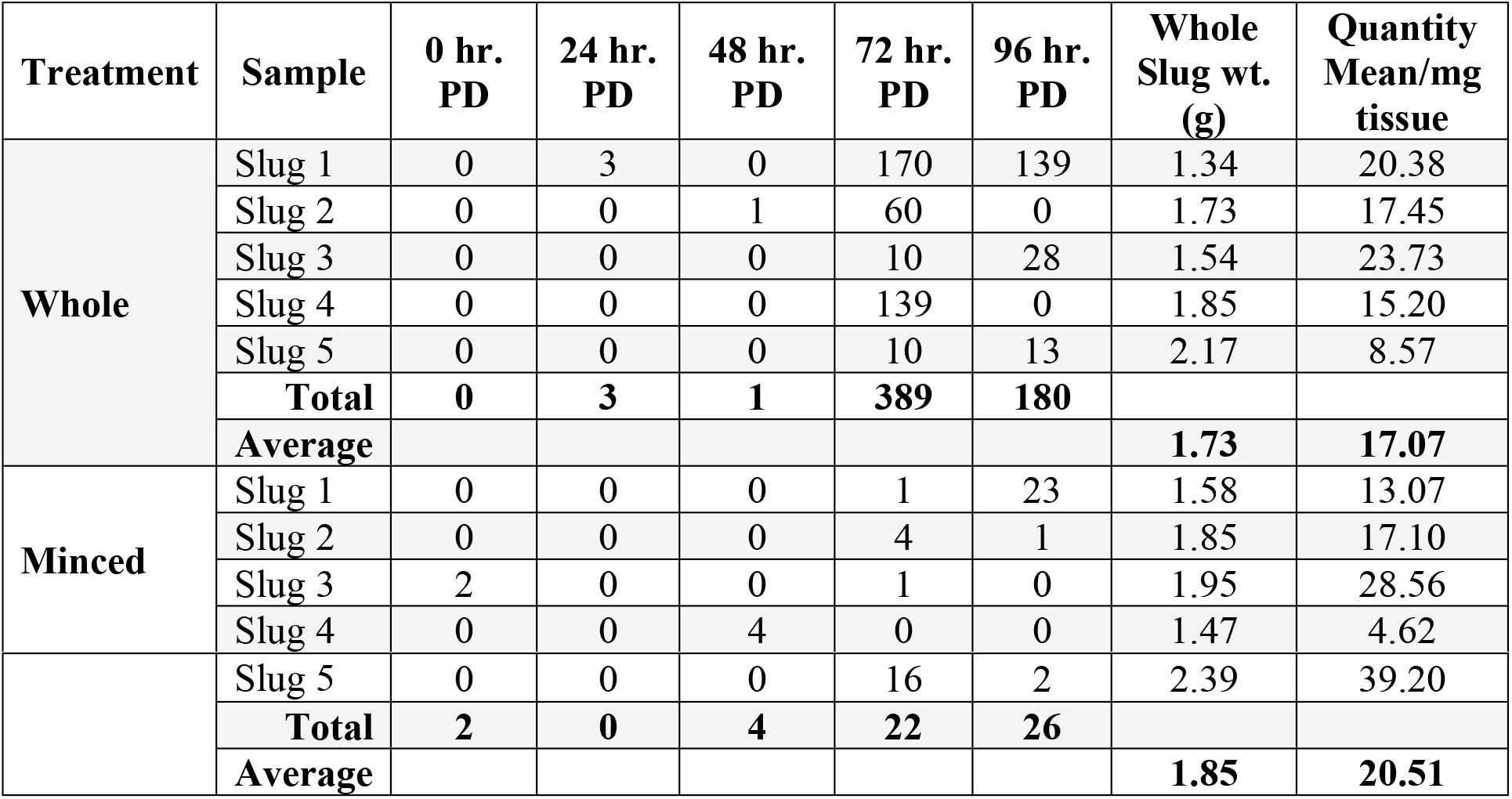
The total number of live larvae shed by the two treatment groups of *P. martensi* at 24, 48, 72, and 96 hours post-drowning. The average weight of the gastropods between treatment groups is not significant (P = 0.586) and the mean quantity of larvae per milligram of tissue estimated by real-time PCR between treatment groups is not significant (P = 0.590), however the numbers shed between treatment groups was significant (P = 0.043). (PD = post drowning).

| Treatment | Sample | 0 hr.<br>PD | 24 hr.<br>PD | 48 hr.<br>PD | 72 hr.<br>PD | 96 hr.<br>PD | Whole<br>Slug wt.<br>(g) | Quantity<br>Mean/mg<br>tissue |
| --- | --- | --- | --- | --- | --- | --- | --- | --- |
| Whole | Slug 1 | 0 | 3 | 0 | 170 | 139 | 1.34 | 20.38 |
|  | Slug 2 | 0 | 0 | 1 | 60 | 0 | 1.73 | 17.45 |
|  | Slug 3 | 0 | 0 | 0 | 10 | 28 | 1.54 | 23.73 |
|  | Slug 4 | 0 | 0 | 0 | 139 | 0 | 1.85 | 15.20 |
|  | Slug 5 | 0 | 0 | 0 | 10 | 13 | 2.17 | 8.57 |
|  | **Total** | **0** | **3** | **1** | **389** | **180** |  |  |
|  | **Average** |  |  |  |  |  | **1.73** | **17.07** |
| Minced | Slug 1 | 0 | 0 | 0 | 1 | 23 | 1.58 | 13.07 |
|  | Slug 2 | 0 | 0 | 0 | 4 | 1 | 1.85 | 17.10 |
|  | Slug 3 | 2 | 0 | 0 | 1 | 0 | 1.95 | 28.56 |
|  | Slug 4 | 0 | 0 | 4 | 0 | 0 | 1.47 | 4.62 |
|  | Slug 5 | 0 | 0 | 0 | 16 | 2 | 2.39 | 39.20 |
|  | **Total** | **2** | **0** | **4** | **22** | **26** |  |  |
|  | **Average** |  |  |  |  |  | **1.85** | **20.51** |

#### Diversified gastropod species

Of the gastropods used in this study, only those that shed larvae or whose tail snips were positive for *A. canotonensis* by qPCR are reported. Two *L. alte* tested positive by qPCR; however, only the minced *L. alte* shed larvae. An unminced *A. fulica* had positive qPCR results but did not shed larvae. Two of the whole *P. martensi* shed larvae and one was positive by qPCR. Coiled, C-shaped, and motile larvae were again observed in samples taken at 48 hours PD. Coiled larvae (L3) emerged into active larvae displaying the S and Q motion, while C-shaped larvae (L2 larvae) were never observed to become motile. Again, the greatest number of larvae found were in bottom samples (95.4%). Low numbers of larvae were shed in the first 48 hours, after which the indicidence of shedding increased, and then began to drop off after 96 hours PD, however, very low numbers of larvae continued to be shed up to 17 days PD.

One qPCR negative *P. martensi* and one positive *L. alte* shed copious amounts of larvae which were observed in petri dishes over time. In addition to the coiled, C-shaped larvae, and several motile larvae in samples taken at 48 hours PD, there was observed in the petri dishes containing the 5 mL samples taken at 24 and 48 hours PD, a gradual emergence of vigorously moving, varied-sized larvae, the numbers of which increased over time, peaking at seven and eleven days after the sample was drawn, with counts of ~ 900 larvae in some dishes (Fig. S1). These larvae were likely emerging from tissue and slime shed by the drowned gastropods. Some of these larvae exhibited the S and Q-movement said to be specific to *A. cantonensis* and when observed by microscopy, these larvae had the clear distinction at the esophagus-intestine junction and a posterior section dense with refractive granules (Fig 4a), indicative of L1 larvae [33]. Real-time PCR of a concentrated sample of several hundred larvae was positive for *A. cantonensis* indicating a presence of this nematode in the sample.

**Fig 4.**
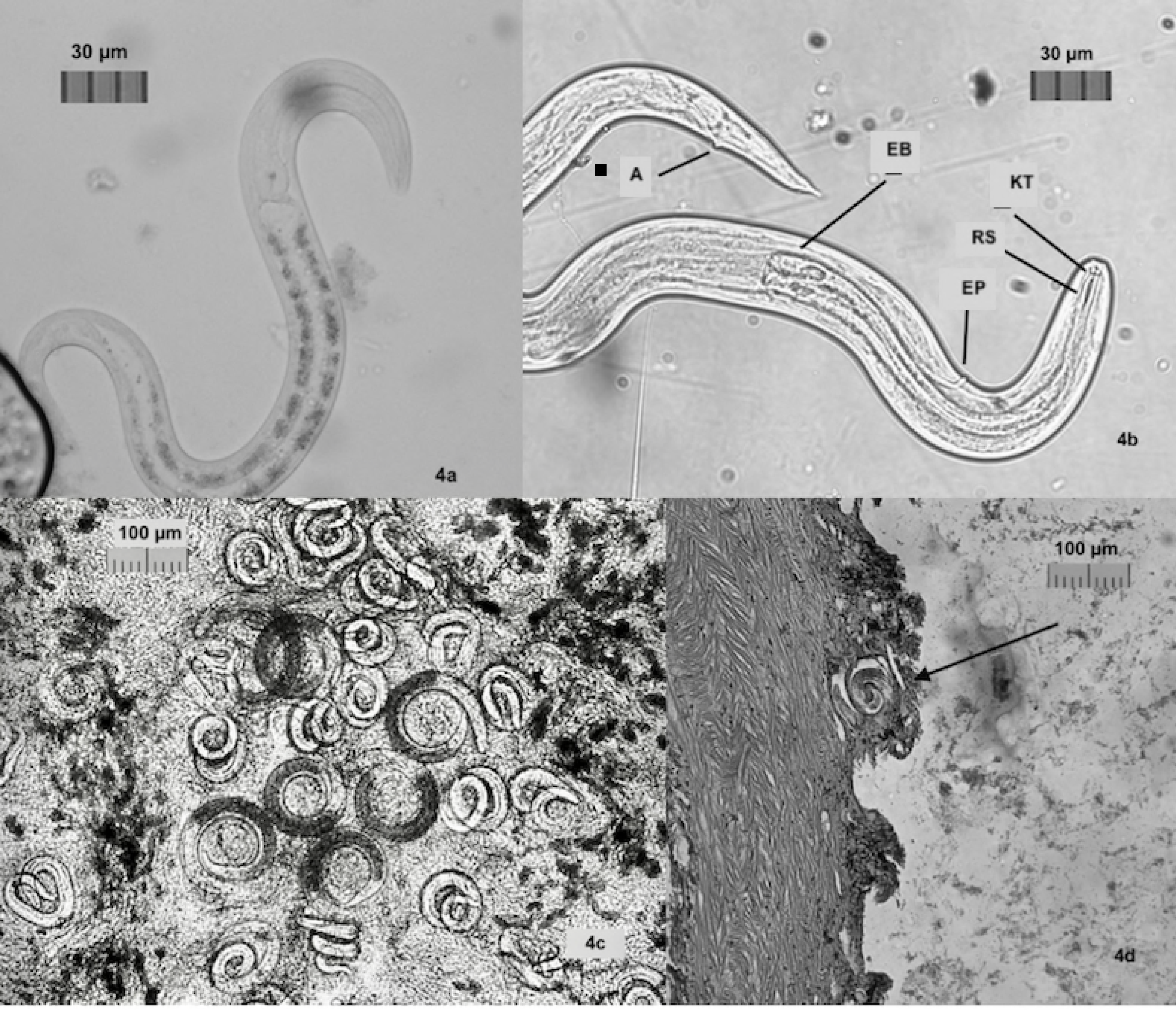
Images of *A. cantonensis*. a. L1 larvae with distinctive junction of esophagus and intestine in the anterior section, posterior section is dense with reflective granules (10μm). b. L3 larvae with knob-like tips (KT) and rod-like structure (RT) in head, clear division of esophagus bulbus (EB), excretory pore (EP), and anus (A). c. Tissue squash from *P. martensi* showing C-shaped L2 larvae with dark interiors. d. Histology section of *P. martensi* showing coiled larvae very close to edge of the body wall.

#### Sieve separation of larval stages and longevity trials

The use of a 20 μm sieve allowed for sorting of varied-sized larvae to determine larval stage and longevity of motile L1 and L3 larvae. Larvae found below the sieve were held alive for 53 days and 56 days respectively, and when a subsample was subjected to a 0.5% HCl/pepsin mix, larvae dissolved. Real-time PCR testing of ~ 250 larvae held for molecular analysis showed positive results for *A. cantonensis.* These appeared to be L1 larvae based on size, sensitivity to acid, and real-time PCR results. Coiled larvae that had emerged into swimming larvae were not able to traverse the sieve and the addition of 10 mL of 0.5% HCl to a subsample did not cause larvae to dissolve. Larvae held in water were active for at least 21 days, with activity decreasing over time. When acid was added to this subsample, the motionless larvae became vigorously active again. Structural characteristics of *A. cantonensis* [33] could clearly be recognized, including the knob-like tips and a rod-like structure in the head, the clear division of the esophagus bulbus, the excretory pore, and the anus (Fig 4b). Real-time PCR of 80 of these active larvae showed positive results for *A. cantonensis.* Based on all of these findings, these appeared to be L3 larvae.

#### *P. martensi* (municipal versus rainwater)

All *P. martensi* with tail snips taken (n = 10) but otherwise left whole were observed to shed larvae at 24 hours and continued to shed larvae up to 96 hours PD. All 10 slugs and shed larval samples were positive for *A. cantonensis* by real-time PCR. All entirely whole (no tail snips taken) *P. martensi* (n = 6) shed larvae at 24 hours and continued to shed larvae up to 72 hours at which point observations were concluded. Shed larvae were again viewed as coiled and C-shaped. One of the whole slugs drowned in rainwater released a clear, mucilaginous mass (presumed to be slime) that contained a count of 328 larvae at day 5 PD.

#### Tissue squashes to screen for presence of larvae

Real-time PCR of all 10 specimens with tail tissue taken showed positive results for *A. cantonensis.* All tissue squashes revealed both coiled and C-shaped larvae. The C-shaped larvae had a clear distinction between the esophagus-intestine line indicative of L2 larvae [33, 34] (Fig 4c).

### Histology

Traditional histological techniques were useful in examining the location of larvae in the *P. martensi* host. Larvae were found throughout the body and were often located near the foot, mantle covering, and very close to the body wall (Fig 4d).

### Catchment Sediment Filter Testing

Live nematodes, including *A. cantonensis* larvae, were able to traverse all sediment filters except that of the 5 μm carbon block filter (Table 2). There were no significant differences found between proportions of post-filter nematodes between replicate runs for the 20 μm (P = 0.999), 10 μm (P = 0.081), and 5 μm spun (P = 0.558) filters. There were no significant differences found between proportions of post-filter nematodes between runs within each filter for the 20 μm (P = 0.180), 10 μm (P = 0.867), and 5 μm spun (P = 0.105) filters. There were no values for the 5 μm carbon block and the 1 μm filter, so no P-values could be generated. A significant difference was found in comparisons between filter type (P =0.0001), with the 10 μm having the highest proportion of post-filter nematodes as compared to the other filters. All positive filtrates contained nematodes with widths greater than the micron size listed for the manufacturer’s nominal filtration rating. Nematodes not recovered during testing, not attributable to the sediment filter, averaged 30 ± 8.5% (mean ± SD), and nematodes not recovered during vacuum filtration alone averaged 22 ± 6.2%. Depending on the filter in the system, water pressure during testing ranged from 25-32 psi; the greatest pressure was seen with the 5 μm carbon block filter. This pressure was comparable to a system with a pressure switch setting of 20-40 psi. Live nematodes, which exhibited S and Q swimming movements associated with *A. cantonensis* larvae [33], were observed in all positive filtrates except the first 1 μm replicate filter. The majority of all pre (88%) and post-filtered (74%) nematodes was within the length range of *A. cantonensis* L3 larvae, which was also true of most filter replicates (Table 2). Post-filter nematodes tested by real-time PCR were positive for *A. cantonensis*, except the third 5 μm polypropylene replicate.

**Table 2.**
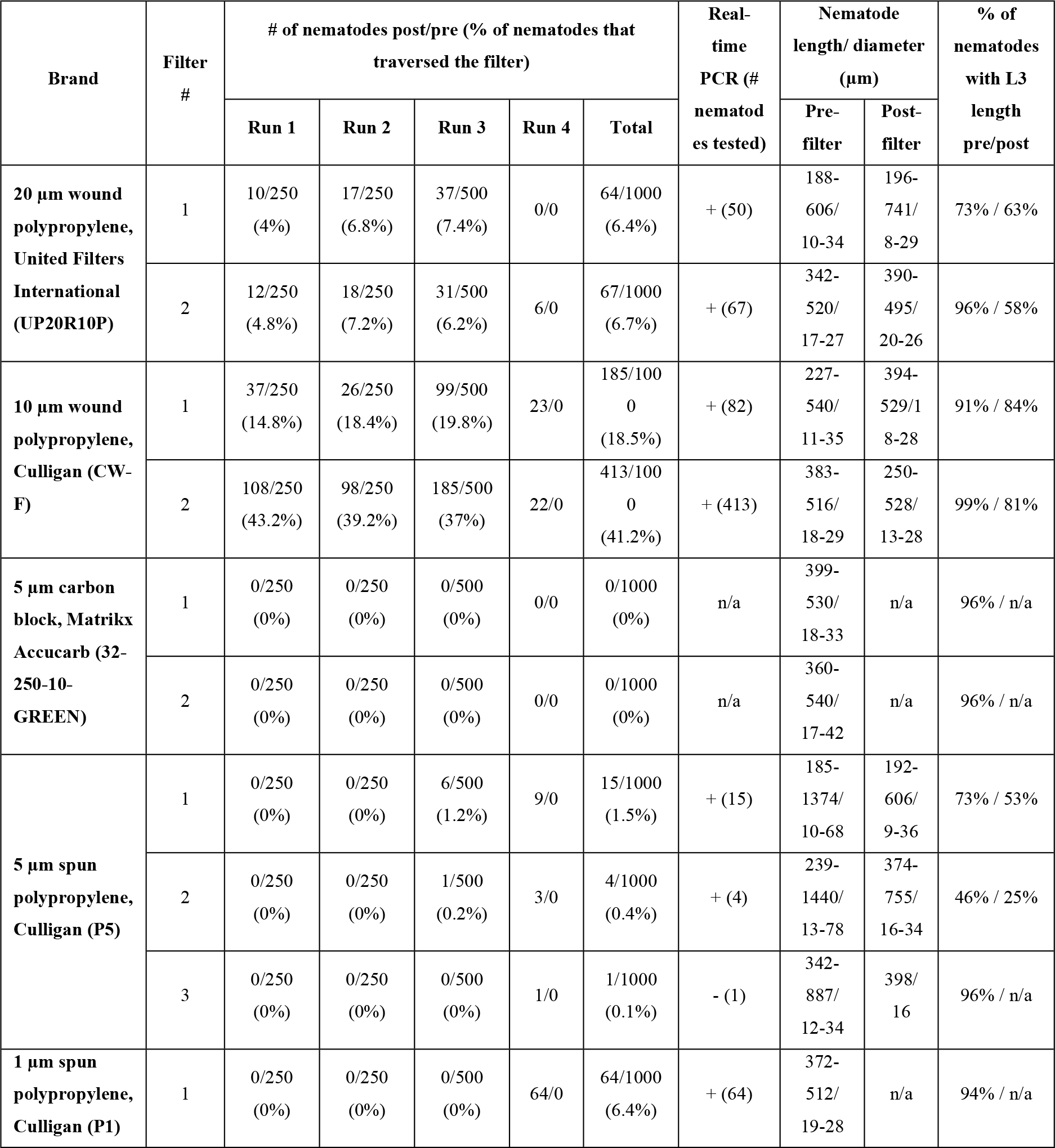
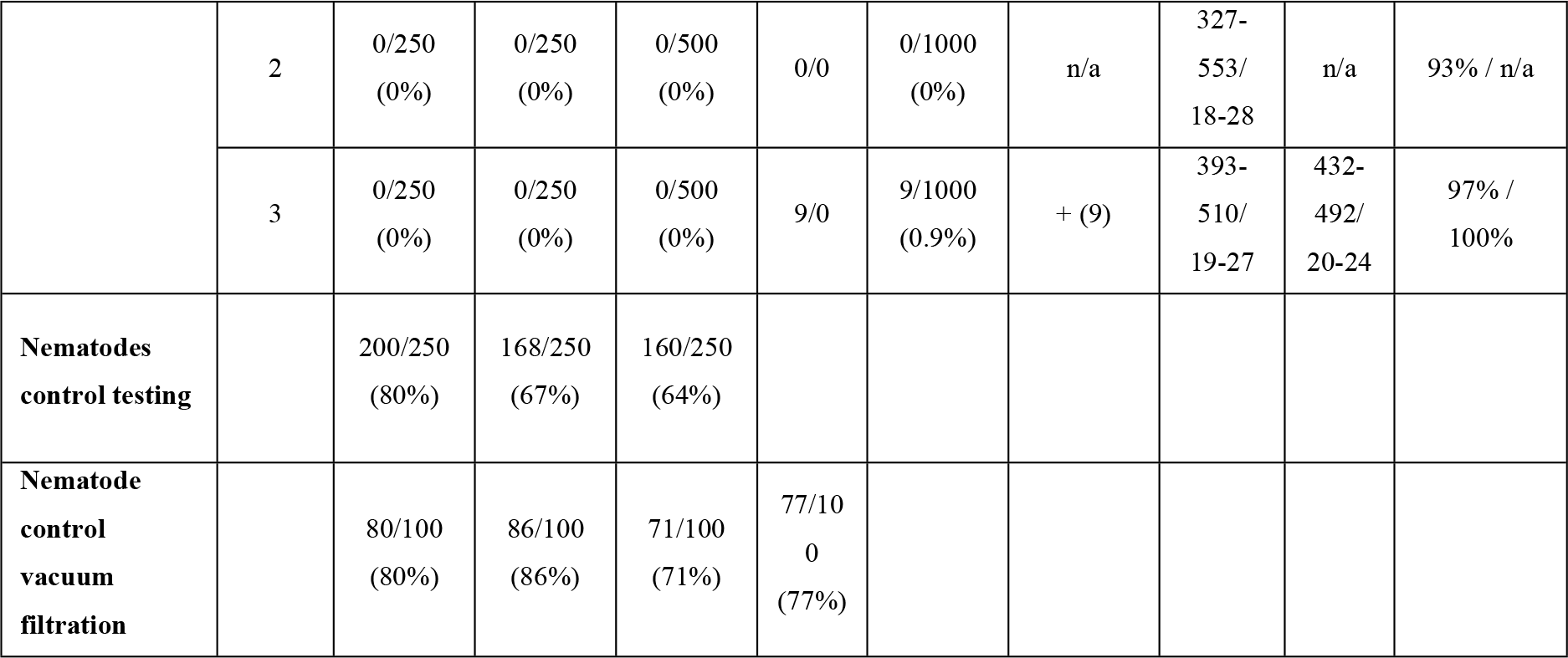
Filters tested, numbers and dimensions of nematodes pre- and post-filtration, qPCR results, including nematode loss controls for the test processes and vacuum filtration.

### Genetic Analysis

Results for qPCR and real-time PCR of samples used in experiments are reported in each respective section above. Results using dilutions of a plasmid as standards and positive controls in qPCR and real-time PCR are reported here. Sequencing and GenBank BLAST analysis verified the plasmid ITS region as an insert (RLW Acan ITS plasmid 18675.11 sequence, 5’ −>3’ TATCATCGCATATCTACTATACGCATGTGACACCTGATTGACAGGAAATCTTAATGA CCC) with 100% sequence match to known *A. cantonensis* ITS sequences (GenBank accession GU587745 to GU587762) [31]. Cycle threshold (C_T_) values of the standards ranged from 15-28 cycles and the plasmids were 15-32 cycles. The standard curve had an R^2^ value of 0.976, a slope of −3.798, a y-intercept of 20.535, and PCR efficiency of 83.354%. The mean quantities of the eight plasmid dilutions ranged from 17.3 to 0.001 larvae per reaction.

## Discussion

These studies substantiate the epidemiological significance of contaminated water as a source of *A. cantonensis* transmission. We have clearly demonstrated the potential for shedding of the infective stage larvae in water from drowned *P. martensi*, a highly efficient, intermediate host, recently introduced to Hawai‘i. In contrast to previous studies, our results demonstrate that undamaged *P. martensi* are capable of shedding several hundred infective stage larvae that can survive in water for several weeks. While current rainwater catchment guidelines state that a 20 μm sediment filter should be sufficient to block the infective stage larvae, our findings show that live, infective-stage larvae were able to traverse 20, 10, 5, and 1μm commercially available wound or spun polypropylene sediment filters.

Rainwater was never observed to contain larvae and real-time PCR results for a water sample showed no presence of *A. cantonensis* DNA. Intact, infected, drowned *P. martensi* shed significantly greater numbers of *A. cantonensis* larvae than minced *P. martensi*, and this finding did not correlate to slug weight or larval burden. Shed larvae included L1, L2 and L3 stage *A. cantonensis*, and in a wet environment, larvae can survive for quite some time; L1 larvae for at least 56 days and L3 up to 21 days. Previously active L3 larvae, which appeared motionless at 21 days PD, were stimulated into activity by exposure to acid; however, additional studies would need to be done to determine if these were infective. Acid destroys L1-L2 larvae; the use of water (as opposed to acid) facilitated the release of these stages and preserved them for study.

The gastropods used in this study were infected in the natural environment and likely harbored other nematode species and/or multiple stages of *A. cantonensis* larvae. As the size of different *A. cantonensis* larval stages is well-documented, a 20 μm sieve was useful for determining larval stages. While feeding L3 larvae to rats is the gold standard for identifying nematode larvae as *A. cantonensis*, funding restraints precluded such studies. Real-time PCR was used to detect the presence of *A. cantonensis* DNA.

In this study, the route through which the larvae exited the drowned slug host was not determined. Histological sectioning of *P. martensi* showed coiled larvae located very close to the body wall, and it is possible larvae may exit the slug via mucus secretions or tissue decomposition. Gastropods can produce mucus for locomotion, to maintain external body moisture, and as a defense mechanism [35], Hyperhydration and differences in somatic pressures may cause the release of mucus, especially in total immersion in water resulting in death [35], Hyperhydration may lead to blood (haemolymph) venting [36] through the pneumostome, which may have caused the release of the L1 larvae. It has been reported within the first 24 hours of infection that larvae may move throughout the slug and may be found in the hemocoel which contains haemolymph [29]. More than 300 live larvae were found 5 days PD in a mucus mass that was exuded from a *P. martensi*, which at 24 hours PD contained only eight visible larvae. The origin of the shed larvae from a drowned slug could the pneumostome, mucus glands, or tissue decomposition.

Tissue squashes may serve as a useful method to observe larvae [4]. Larvae, both coiled and C-shaped, were clearly visible in the tissue squashes without staining. While molecular analysis is still necessary for confirmation of *A. cantonensis*, the tissue squash technique could work quite well for screening gastropods for infection. Slugs found in commercially bought salads have been brought to our lab for testing. In high infection areas such as Hawai‘i, a tissue squash may provide the visual evidence needed to immediately and prophylactically administer anthelmintic drugs in the case of human exposure.

L1 larvae passed through a 20μm metal sieve, but L3 larvae were unable to traverse the sieve. While no L3 larvae were able to migrate through the 20μm metal sieve, the infective stage L3 larvae are capable of burrowing through the intestinal wall, and while they may not be able to burrow through metal, they may be capable of migrating through a non-metal filter. We evaluated the ability of nematodes isolated from wild-caught *P. martensi* to traverse five different types of sediment filters commonly used in household catchment systems. While live nematodes were able to traverse all filters except the 5 μm carbon-block filter, all filters significantly reduced the number of nematodes introduced to the system. We believe the structural design and differences in construction of individual filters are important variables in determining if nematodes are able to traverse the filters tested. Similar to the metal sieve, the carbon-block filter is the only filter tested that is made of inflexible material (100% coconut shell carbon) and possesses rubber seals on each end. Nematodes could not go around the carbon filter swept by water currents or swimming while the system was off, nor could they burrow or swim through the carbon filter while the system was off. While only two carbon filters of one brand were tested, future research should particularly focus on other brands and sizes of carbon block filters, with larger sample sizes, for testing effectiveness for blocking nematodes. Structural design and construction differences also likely explain the finding of more nematodes in the 10 μm filtrate than the 20 μm filtrate, as the 10 μm filter had thinner strings that were notably more loosely wound compared to the 20 μm strings. There was even a clear difference in string tightness between the two 10 μm filters tested, which likely caused the large but not significant differences in the proportion of post-filter nematodes between each filter replicate. The other filters showed no significant differences in proportions of post-filter nematodes between filter replicates, indicating the construction of some filters can be consistent and produce reliable results. The diameters of the nematodes found in the filtrates were greater than the manufacturer’s listed micron size. Despite all of these findings, to the best of our detection capabilities, it seems the 20 μm, 5 μm spun, and 1μm filters performed as stated by the manufacturer nominal ratings which reduce >85% of particles with the listed micron size. The 5 pm carbon block outperformed these standards, while neither replicate of the 10 μm filter met this standard. Most commercially available catchment filters are not rated with ‘absolute’ microns due to their structure. We suspect that there will be some flex in the filter micron size based on the structure and material of the filter, thus most filters are considered ‘nominal’.

While other nematode species were likely present in the catchment filter tests, since some length and width measurements were larger than known for *A. cantonensis* L1-L3 larvae, the majority of pre- and post-filtered nematodes were within the length range of *A. cantonensis* L3 larvae (Table 2). Moreover, many live post-filter nematodes exhibited swimming S and Q-movement patterns indicative of *A. cantonensis* larvae [33]. Real-time PCR results of filtrates with more than one nematode were all positive, and the lack of *A. cantonensis* detection of the one nematode from the third replicate of the 5 μm spun filter was likely due to DNA concentrations of the extraction being below the sensitivity of the real-time PCR assay. Together, the nematode sizes, swimming patterns, and genetic analyses indicate that live *A. cantonensis* larvae were among those that traversed the filters.

This study is especially important for Hawai‘i because of the widespread, unregulated use of rainwater catchment systems for household water supplies. While all of the filters, except the 10 pm, performed at or above the manufacturer’s ratings, clearly there is misplaced trust by contractors, vendors, and homeowners in the effectiveness of sediment filters to completely block larger parasites like *A. cantonensis.* Outreach should be done to educate rainwater catchment users in high infection areas about the meaning of nominal filter ratings. Additional research should also verify that *A. cantonensis* cannot traverse filters with absolute filter ratings.

Given the limits of our detection capabilities based on the methods used, as well as the limited sample size of sediment filters tested, homeowners should be cautious in relying on the same model of filters used in their own catchment systems to perform identically to the results reported here. This study was conducted in an isolated, clean, laboratory environment which may be quite different from a homeowner’s catchment system regarding reservoir debris, water pump strength, and overall system maintenance. It is unknown if the buildup of organic debris on sediment filters affects either the viability of *A. cantonensis* larvae or the filtration capabilities throughout the course of the filter’s life. It is also unknown if a water pump that generates higher pressure on the sediment filter would produce different results. Moreover, homeowners should not extrapolate the results of this study to other brands of filters constructed with similar materials or micron ratings.

Disinfection treatments might play a crucial role in protecting households. Chlorine has been shown to kill microorganisms in water, including the nematode *Angiostrongylus costaricensis* [37], and has been recommended to treat catchment reservoirs [21]. Bleach, however, is not FDA approved for water treatment by private citizens and it reacts in water with natural organic matter to produce toxic halogenated volatile organic compounds called trihalomethanes [38]. NucShort-wave ultraviolet light (UVC, 254 nm) is also a widely endorsed disinfection method, however eukaryotic organisms like nematodes possess the ability to repair nuclear DNA damage from UV radiation [39]. These findings showcase that even ‘properly designed’ rainwater catchment systems may leave users exposed to *A. cantonensis*. Novel ideas may be needed to address this problem, not only for Hawai‘i residents but also for rainwater catchment users across the globe in regions where parasitic nematodes are endemic.

It is feasible that parasitized gastropods or paratenic flatworms that perish and decompose in wet field crops, or those that are smashed in roadway puddles or drowned in small bodies of water, may release infective stage nematodes capable of disease transmission. Instances of human infection have been recorded in Texas following a flood event [40, 41], and while transmission via ingestion is generally considered the primary source of infection, a mouse study shows the potential for *A. cantonensis* transmission through oral, intraperitoneal, sub-cutaneous, lacerated and unabraded skin, anal, vaginal, conjunctival mucosal tissue, and foot pad, but not tail penetration [42]. Intraperitoneal and subcutaneous injection resulted in recovery of more worms than from oral intubation. The study concluded that skin or mucosa contacts with L3 *A. cantonensis* larvae may be a cause of angiostrongyliasis/neuroangiostrongyliasis in the natural environment.

Likewise, beverages left outside uncovered could become a repository for a wandering gastropod and a source of disease transmission. In Hawai’i in 2017 there were two confirmed cases and four probable cases of neuroangiostrongyliasis resulting from consumption of homemade kava, a traditional drink common to the western Pacific islands. The kava had been left in an uncovered bucket and a slug was found in the bottom of the container holding the beverage after the kava was consumed [43]. Recently, a case of neuroangiostrongyliasis was reported in Hawai‘i and the victim reported drinking from a garden hose [44]. Because residents report finding *P. martensi* in hoses, this should also be considered as a potential pathway for disease transmission.

## Conclusion

It is not improbable that the widespread use of rainwater catchment as household and agricultural water supplies may play a role in the high number of cases of neuroangiostrongyliasis originating from East Hawai‘i Island, and this public health concern should be thoroughly investigated. As rainwater catchment use is unregulated in Hawai‘i, systems may potentially be a cause of not only rat lungworm disease, but also other diseases found in Hawai‘i, such as leptospirosis, giardia, salmonella, and *Escherichia coli* infections, all of which can be transmitted through water. It is important to understand that the Puna District of Hawai‘i Island is one of the fastest growing areas in the entire U.S. due to availability of affordable land and large subdivisions that contain many lots available for sale. Without research and education-related intervention, case numbers of neuroangiostrongyliasis may continue to rise in Hawai‘i. It is essential for epidemiologists to consider and investigate rainwater catchment systems as potential pathways for *A. cantonensis* transmission.

## ACKNOWLEDGEMENTS

We thank Hawaii Catchment Co., Keaau, and Keaau Ace Hardware, Hawaii for providing filters and equipment needed to complete this study. Many thanks to Ann Kobsa, Lyn Howe and Geoff Rauch for the generous supply of gastropods, and to Deborah Beirne and the Department of Biology for the loan of a dissecting microscope for these studies. To Michael Severino we extend our sincere gratitude for your assistance with the molecular analysis of gastropod tissue. We thank Dr. Alex Hanlon for assistance with the statistical analyses, and we are very grateful to Roxana Myers and Cathy Mello from the USDA Pacific Basic Agricultural Research Center, and Dr. Adler Dillman from Stanford University, for their assistance with nematode identification. Many thanks to Patricia Macomber for her insights regarding rainwater catchment use on Hawai‘i Island and Kirsten Snook for review of the manuscript. Also, a very special thank you to Dr. Marta DeMaintenon at the University of Hawai‘i at Hilo for her aid and expertise in the histological preparation of gastropods.

## Supporting information

**S1 Fig. Many and various sized larvae.** Larvae appeared in samples taken at 24 and 48-hours PD from drowned gastropods. Low numbers were visible in the dish upon initial inspection, however numbers of larvae in the petri increased over time, peaking at days seven and eleven. **https://www.youtube.com/watch?v=CkLCBeqFRW4**

